# Dose-dependent effect of ketamine on functional connectivity in mice

**DOI:** 10.64898/2026.09.03.749142

**Authors:** Tomokazu Tsurugizawa, Aya Takemura

## Abstract

A single injection of a subanesthetic dose of ketamine, an NMDA receptor antagonist, induces schizophrenia-like behavior in mice. Previous studies investigated the effect of single injection of ketamine on behavior and c-Fos expression in mice dose-dependently. However, there has not been functional magnetic resonance imaging (fMRI) study to investigate the effect of ketamine in mice. Here, we used awake fMRI and investigated the dose-dependent effect of single injection of ketamine in awaked mice. The lower-dose (5 mg/kg) ketamine increased functional connectivity (FC) with the thalamus and the hypothalamic nuclei. Higher-dose (50 mg/kg) ketamine increased widespread FC with cerebral cortex as well as hippocampus and thalamic nuclei. Graph theory analysis reveals that several regions of hypothalamus and cerebral cortex are essential for the network change by ketamine injection. These results indicate that sub-anesthetic ketamine dose-dependently alters the brain and indues the abnormal behavior in mice.

## 1. Introduction

A single injection of a subanesthetic dose of ketamine, an NMDA receptor antagonist, induces schizophrenia-like behavior in mice [1]. Ketamine injection affects this behavior in a dose-dependent manner. The injection of more than 50 mg/kg ketamine impairs working memory in the Y-maze and sensorimotor gating, as assessed by the inhibition of the acoustic startle response in mice [1]. It also induced hyperlocomotion during an open field test. Similarly, at a lower dose (20 mg/kg), the behavioral changes were mild and transient. In contrast, a higher dose (50 mg/kg) induced marked stereotypical behaviors, including head weaving, circling, ataxia, and hyperlocomotion [2]. The injection of lower dose ketamine (3–30 mg/kg) during mid-adolescence promoted food intake and body weight in female mice with activity-based anorexia [3].

A previous study used calcium imaging to investigate the effect of anesthesia (ketamine/xylazine) on the synchronization of slow calcium oscillations in awake mice [4]. The injection of 10 mg/kg of ketamine in rats induced increased transcripts of NUCB2, GPR173, and POMC [5]. The acute injection of ketamine (20 mg/kg) increased the number of c-Fos-immunoreactive neurons in the dentate gyrus of the hippocampus, field CA1 of the hippocampus, dorsal subiculum, ventral subiculum, ventral subiculum, central amygdaloid nucleus, and basolateral amygdaloid nucleus [6,7]. Ketamine injection induces c-Fos expression in a dose-dependent manner [2]. Higher doses of ketamine (50 mg/kg) induces the c-Fos-immunoreactive neurons in the cingulate cortex and the retrosplenial cortex but this c-Fos induction is weak in lower dose of ketamine (20 mg/kg). We therefore hypothesized that ketamine affects the brain activity and alters behavior in a dose-dependent manner in mice.

To elucidate the effects of ketamine the neuronal activity before and after ketamine injection on the entire brain should be compared. Functional connectivity (FC), which reflects the synchronization of fluctuations in blood oxygenation level-dependent signals among anatomically separated brain regions and reflects the functional network in the brain, can be estimated using functional magnetic resonance imaging (fMRI) [8]. A clinical study revealed alterations in FC by ketamine injection in healthy volunteers and patients with schizophrenia and acute ketamine injection increased PFC-HC connectivity in healthy volunteers [9]. Although clinical study reveals that the effect of ketamine on FC directly, it is difficult to investigate dose-dependent manner more deeply. The animal model is suitable to investigate the behavior and neuronal activity in detail. The recent development of mouse fMRI enables awake fMRI to be performed [10,11]. However, in contrast to clinical fMRI, mouse fMRI shows local activation following ketamine injection (1 mg/kg) and no studies have shown altered FC [12]. Previously, we induced anesthesia in awake mice in MRI and successfully demonstrated the effects of light anesthesia induction in conscious mice [13]. The advantage of this technique is that it allows for direct comparison between the anesthetized and awake states in the same mice. Here, we used this method to elucidate the dose-dependent effects of ketamine in awake mice. We also used the graph theory to investigate the key brain regions for the altered FC.

## 2. Method

### 2.1 Animals and surgery

Ten male C576J/BL mice (18–25 g, 10–15 weeks old at the start of surgery) were used. Male mice were used to avoid the effects of the estrous cycle in female mice. The sample size was determined based on previous studies [13,14]. All animal procedures used in the present study were approved by the Institutional Animal Care and Use Committee of the National Institute of Advanced Industrial Science and Technology. All methods used in this study were performed in accordance with the relevant guidelines and regulations.

### 2.2 Surgery for awake fMRI

We performed surgery on the mice under isoflurane anesthesia (1.5%–2.0% with air) for the awake fMRI [15]. The skin on the heads of the mice was cut longitudinally and the skull surface was polished using saline and etchant gel with gauze (Super-Bond C&B, Sun Medical Company, Ltd., Shiga, Japan). The blood was carefully removed. A peek head pole (0.5 mm diameter), which was used for head fixation without pain during the imaging session, was attached to the skull vertically with resin cement (Super-Bond C&B, Sun Medical Company, Ltd., Shiga, Japan). To avoid susceptibility artifacts, it is important to exclude air and blood between the skull and dental cement when dental cement is mounted on the skull. Following surgery, the mice were allowed to recover for more than 1 week.

### 2.3 fMRI acclimation training

Following recovery from surgery, we confirmed that the mice did not urinate or jump from cupped hands during handling before acclimation training. The detailed setup of the awake fMRI has been described in previous studies [10,15]. The mice were placed on a head positioner by fixing the head pole. To reduce possible stress induced by scanning noise, we used earplugs designed to fit the mouse’s ears (a photograph is shown in Tsurugizawa et al., 2021). To minimize restraint-induced stress, the body was gently wrapped in tissue to allow it to move freely within the tissue cover. The mice were trained for 4 days to become accustomed to the fMRI conditions. A pseudo-MRI system was used during the first 2 days. The mice were anesthetized with 1.5% isoflurane for maintenance and positioned on the pseudo-MRI apparatus. The mice remained in the pseudo-MRI apparatus for 30 min on the first and second days. For the next 2 days, the mice were placed on an MRI bed with a head fixation system using the same methods as in the fMRI experiment (see the awake fMRI section for details regarding the awake resting-state fMRI procedure). Throughout the training period with the MRI system, the respiratory rate was maintained within the normal range using an MR-compatible monitoring system (model 1025; SA Instruments, Stony Brook, NY, USA).

### 2.4 Awake and anesthetized fMRI

All MRI experiments were performed using a Bruker 9.4 T MRI system with a homemade awake fMRI coil (Bruker BioSpin, Ettlingen, Germany) (Fig. 1A). The mice were anesthetized with 3% isoflurane in air for induction, and the isoflurane dose was rapidly changed to less than 1.0% with air for maintenance during head fixation and adjustment of magnetic homogeneity, shimming, and reference pulse. A schematic of the experimental paradigm is shown in Fig. 1B. Isoflurane was administered for no more than 10 min during the adjustment. Anesthesia was stopped, and a structural image scan was started. The mice were awakened during the structural imaging scan. We confirmed that the respiratory rate increased around 150 breaths/min during the recovery period. A previous study showed that neuronal activity recovered to that of a conscious state 10 min after cessation of Iso delivery [10]. The fMRI acquisition was initiated at this point. A schematic paradigm of the MRI protocol is illustrated in Fig. 1B. When the mice were awakened from anesthesia after the recovery period, awake resting-state fMRI was performed. Once the awake fMRI was completed, lower-dose (5 mg/kg body weight) or higher-dose ketamine (50 mg/kg body weight) was injected subcutaneously through the tube via the infusion cannula, which was inserted under the skin on the animal’s back. Resting-state fMRI scanning was started again 5 min after the ketamine injection. For the crossover test, five mice received higher-dose ketamine and then received lower-dose ketamine after 3–5 days, and vice versa for the others. The resting state fMRI images were acquired using a T2*-weighted multi-slice gradient-echo EPI sequence with the following parameters: TR/TE = 2000/18 ms, spatial resolution = 170 × 170 × 700 µm^3^/voxel, matrix size = 88 × 88, 16 slices, and 450 volumes (total: 15 min). For normalization in image processing, a structural image with the same field of view as the fMRI was acquired using T2-weighted multi-slice rapid acquisition with relaxation enhancement (RARE), with the following parameters: TR = 4,000 ms, effective TE = 30 ms, spatial resolution = 117 × 117 × 600 µm^3^/voxel, RARE factor = 8, and 5 averages.

**Figure 1.**
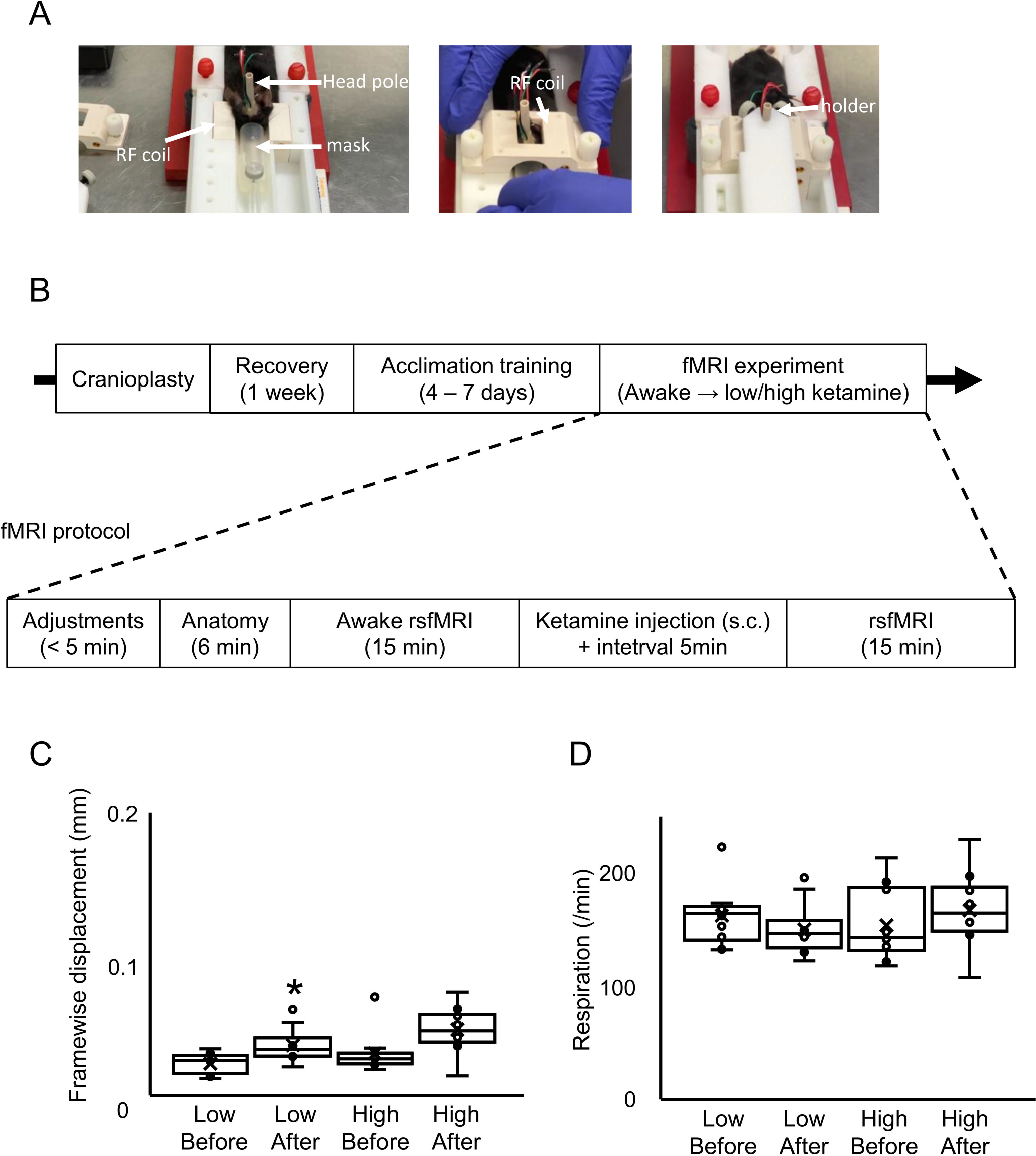
Setup of awake fMRI and schematic figure of the experimental paradigm (A) Photos of awake fMRI setup. The split-type radiofrequency (RF) volume coil, mouse bed, and fixation system of head post. were designed for awake fMRI. (B) Schedule of MRI experiment. (C) framewise displacement and (D) respiratory rate in each condition. *p<0.05, Bonferroni-corrected.

### 2.5 Image preprocessing

The statistical parametric mapping SPM12 software (Wellcome Trust Centre for Neuroimaging, UK) was used to analyze the fMRI data and to perform preprocessing steps, including slice timing correction, motion correction by realignment, normalization, and smoothing with a Gaussian filter (0.6 × 0.6 × 0.6 mm^3^/voxel of the half-width at half-maximum). Before preprocessing, template images coregistered with the Allen Mouse Brain Atlas were obtained (http://atlas.brain-map.org/). The functional and structural images were normalized to these template images. Framewise displacement (FD) was used to verify head motion [16]. The FD was calculated using six motion parameters (three each for translation and three for rotation) from the realignment in SPM. It was confirmed that the head motion for all subjects met the following criteria: (1) the mean FD averaged for all time points during the scan was less than 0.05 mm and (2) the FDs at all time points were less than 0.03 mm. The linear trend in the fMRI dataset time course was removed voxel by voxel using the CONN toolbox (https://web.conn-toolbox.org/) [17]. The mean signals in the ventricles and white matter, and six motion parameters of an object (translational and rotational motions) were regressed from the time series of each voxel to reduce the contribution of physiological noise, such as respiration and head movement.

### 2.6 FC analysis

The preprocessed fMRI data were then detrended, and slow periodic fluctuations were extracted using a bandpass filter (0.01–0.08 Hz). A total of 116 regions of interest (ROIs; 58 per hemisphere) were delineated based on the template images, with reference to the Allen Mouse Brain Atlas (https://scalablebrainatlas.incf.org/mouse/ABA12). The correlation coefficients between two ROIs were calculated using the CONN toolbox. The statistical significance of the ROI–ROI connectivity was assessed by a paired t-test or two-sample t-test with a threshold of p < 0.05 (Network-Based Statistic (NBS)-corrected), using the NBS toolbox (https://www.nitrc.org/projects/nbs/). The connections with a Cohen’s d value exceeding 0.7 (minimum meaningful effect size) were further assessed for NBS. Subnetworks were then identified in the set of suprathreshold connections. The size of each subnetwork was measured in terms of the number of connections that it contained. Permutation testing was used to compute a family-wise error-corrected p-value for each subnetwork. Specifically, 5,000 permutations were generated, in which group labels were randomly permuted, and the size of the largest subnetwork was recorded for each permutation. Subnetworks deemed significant by permutation testing (p < 0.05, family-wise error-corrected) were identified as significant.

### 2.7 Graph theory calculations

Graph theory is advantageous for analyzing complex systems such as FC. We applied graph theory for analyzing the FC using the Brain Connectivity Toolbox [18]. Pairwise connections were reorganized into symmetrical graphs containing 116 nodes and 6,670 edges. The graph edge weights were normalized to a maximum value of 1 (while retaining their normal distribution) and negative connections were set to 0. To avoid any bias in the calculated network measures owing to inter-subject variations in the edge densities, we applied a threshold that maintained an equal edge density across subjects. The graph theoretical measures were calculated across a range of network densities (10%–30%) to reduce threshold selection bias. The area under the curve was computed for each network measure and used for group comparisons by means of the Student’s t-test. Following the density threshold, the connectivity matrices were binarized into graph spaces G such that G_ij_ = 1 indicates the existence of a "connection" between any nodes *i* and *j*. The characteristic path length (L) was calculated as the lowest average number of edges between any pair of nodes *i* and *j* in graph G. We analyzed the mean clustering coefficient (MCC), which is an index of the average number of connected neighbor pairs *j k* of a node *i* being connected to one another [19]. The global efficiency (Eglob) was determined as the average inverse of L. Using these parameters, we calculated the following network measures [20]. The local efficiency (Eloc) describes how well information can be transmitted in the immediate surrounding of node *i* and provides a basis for effective segregated information processing in the network. The degree is calculated as the total nodal connections. The node strength is the sum of all edge weights connected to node *i* in a weighted directed or undirected graph. The participation coefficient is detected if the edges of node *i* tend to be clustered into one module of the network or link different specialized modules.

## 3. Results

### 3.1 Dose-dependent alteration of FC by ketamine injection

The FD significantly increased following lower- dose ketamine injection (p < 0.05, Bonferroni-corrected)and that did not significantly changed by higher-dose ketamine injection (Fig. 1C). Importantly, all FDs are within the similar range of mouse resting state fMRI study in other institutes [21]. The respiratory rate was not changed following ketamine injection (p > 0.05, Bonferroni-corrected) (Fig. 1D). This seems to be consistent with the previous studies showing the increased voluntary activity by ketamine injection [1]. We then investigated the effects of ketamine injection on the FC (Supplementary Fig. 1). To provide whole-brain characterization of the FC, the entire brain was parcellated into 116 broad regions and the Pearson correlation coefficient was calculated between pairs of regionally averaged fMRI time series to yield a 116 × 116 connectivity matrix for each mouse. The injection of lower-dose ketamine significantly increased the FC among the hypothalamus and thalamus bilaterally (Figs. 2A and 2B). In contrast to lower-dose ketamine, higher-dose ketamine significantly increased the FC, including that in the cortex (Figs. 2D and 2E). The 3D figures shows that lower-dose ketamine increased the FC in the ventral and medial parts of the brain (Fig. 2C). Higher-dose ketamine increased widespread connectivity, including in the cerebral cortex (Fig. 2F). The averaged connectivity strength in positive functional connectivity significantly increased by lower-dose ketamine injection (p < 0.05, Bonferroni-corrected) (Supplementary Fig. 2). The injection of higher-dose ketamine increased the averaged connectivity strength but it was not statistically significant.

**Figure 2.**
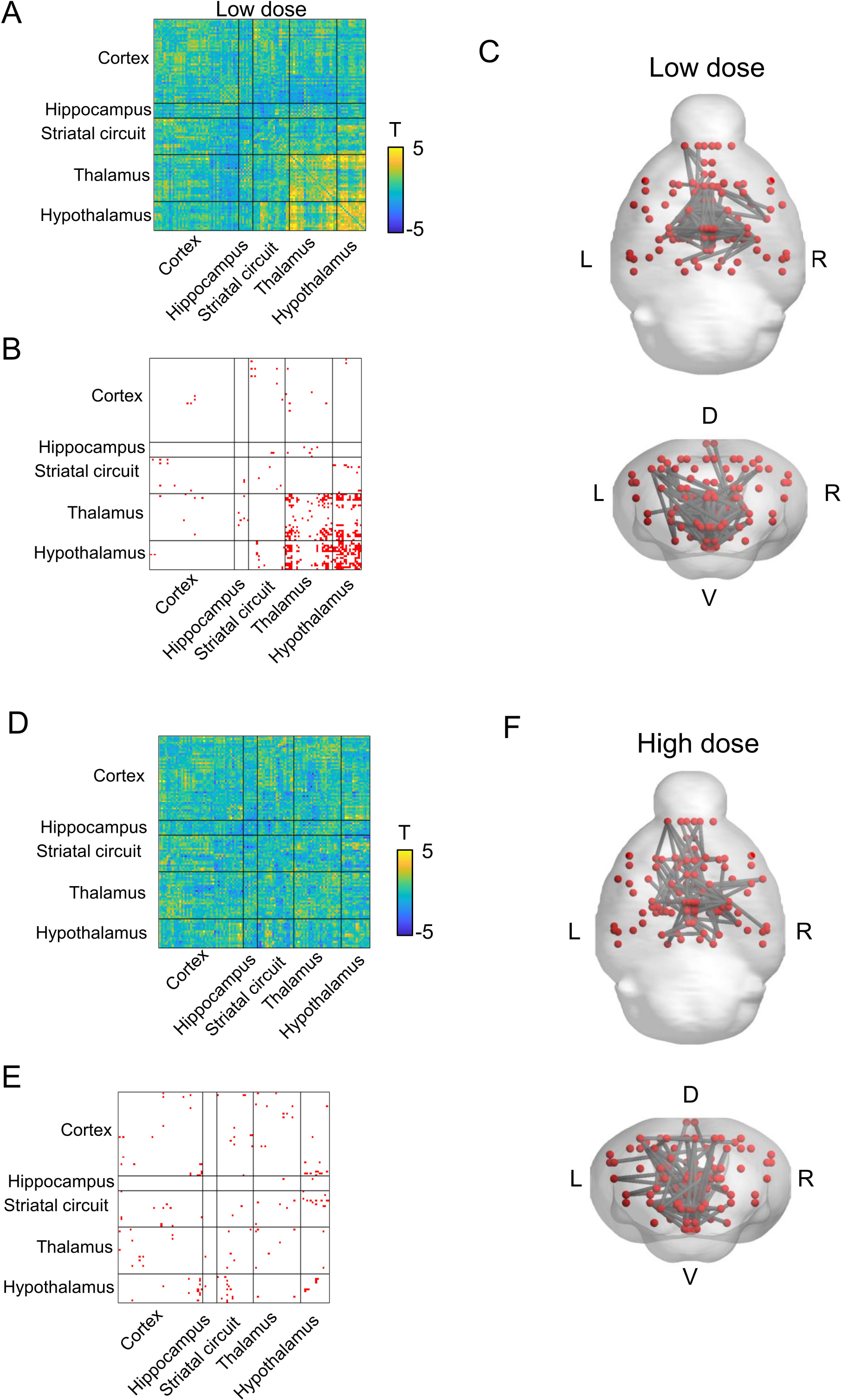
Significant changes of FC following ketamine injection. (A) T-values and (B) significant alteration of Fc following lower-dose ketamine injection (p < 0.05, NBS-corrected paired t-test, n = 10, df = 9). Color bar indicates the T-values. (C) schematic figure that shows the increased FC following lower-dose ketamine injection. (D) T-values and (E) significant alteration of Fc following higher-dose ketamine injection (p < 0.05, NBS-corrected paired t-test, n = 10, df = 9). Color bar indicates the T-values. (C) schematic figure that shows the increased FC following higher-dose ketamine injection.

The significantly increased FC strength with each seed ROI is shown in Fig. 3. The FC strength of a few connections with the orbital cortex, motor cortex, and striatum significantly increased following the injection of lower-dose ketamine (Fig. 3A). Particularly, the significantly increased FC strength was observed in many connections with subregions of the hippocampus, thalamic and hypothalamic nuclei (Fig. 3A). In contrast to the lower-dose group, higher-dose ketamine significantly increased the FC strength in several connections with the orbital cortex and motor cortex (Fig. 3B). Remarkably, the number of the connection showing significant increase of FC strength by higher-dose ketamine with the hippocampus, thalamus, and hypothalamic nuclei was smaller than that by lower-dose ketamine.

**Figure 3.**
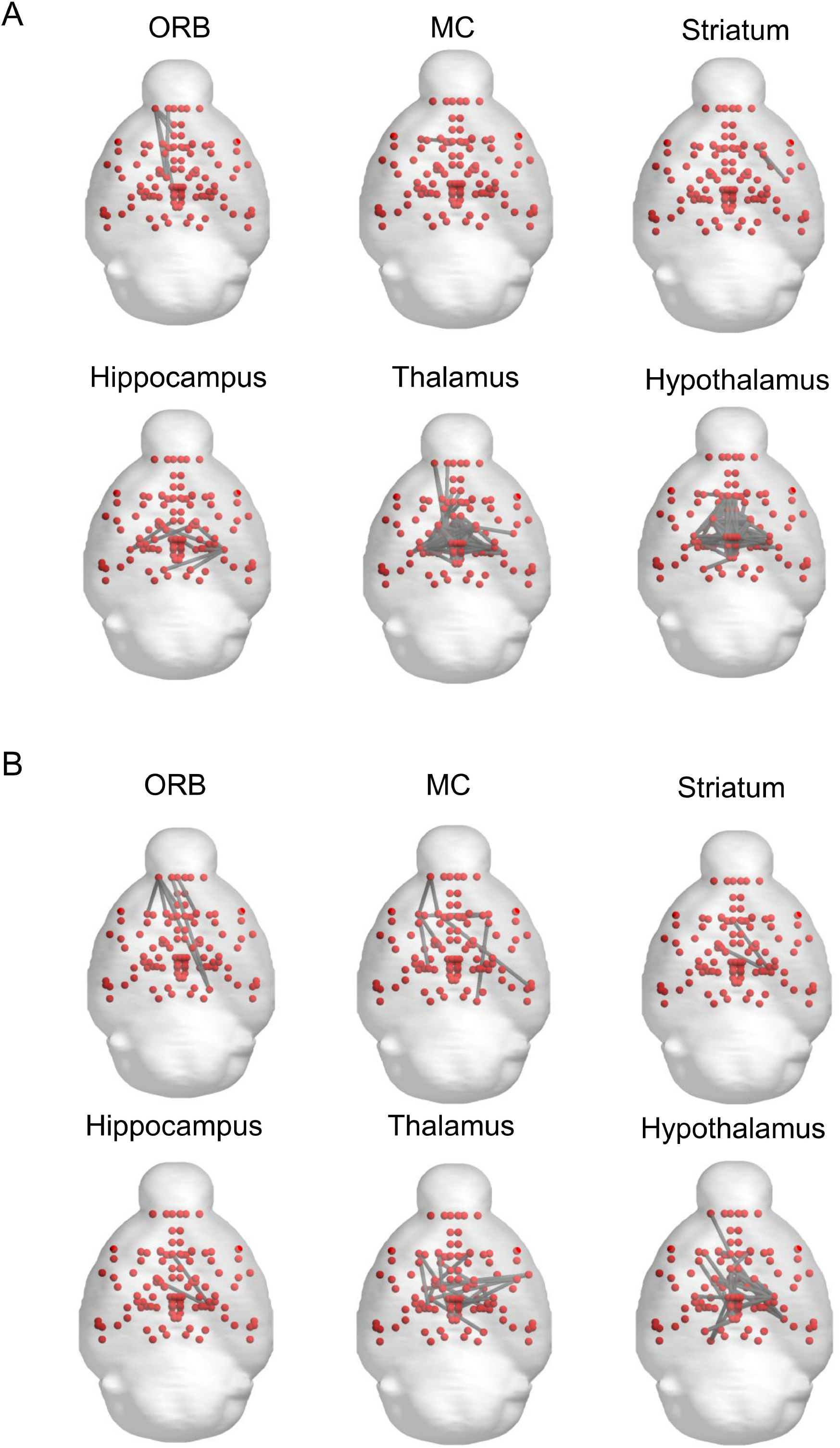
Increased seed-based FC following ketamine injection. (A) Increased seed-based FC with orbital cortex (ORB), primary motor cortex (MC), striatum, hippocampus, thalamus, and the hypothalamus following lower-dose ketamine injection. (B) Increased seed-based FC with orbital cortex (ORB), primary motor cortex (MC), striatum, hippocampus, thalamus, and the hypothalamus following higher-dose ketamine injection. p < 0.05, NBS-corrected paired t-test, n = 10, df = 9.

### 3.2 Key nodes of widespread reduction of static FC under anesthesia

We then attempted to identify key regions involved in the alteration of FC following ketamine injection. The effect size (Cohen’s d) of each connection was calculated from the t-values, and the effect sizes of all connections with each node were summarized. Extensive augmentation in connectivity strength according to effect size with lower-dose ketamine were observed, mainly in the thalamic nuclei (ventral medial nucleus (VM), ventral anterior-lateral complex (VAL), centromedial nucleus (CM), anterior-medial nucleus (AM), mediodorsal nucleus (MD), anterior-ventral nucleus (AV), ventral posteromedial nucleus (VPM), and ventral posterolateral nucleus (VPL)) and hypothalamus (posterior nucleus, lateral hypothalamic area, dorsal part of posterior nucleus (dPH), lateral hypothalamus (LH), medial preoptic area (mPOA), dorsomedial nucleus (DMH), ventromedial nucleus (VMH), and arcuate nucleus (ARH)) (Fig. 4A). In contrast to lower-dose ketamine, higher-dose ketamine strengthened the different nodes, such as the cerebral cortex (primary and secondary motor cortex (M1 and M2), low-limb of secondary somatosensory cortex (SS2-ll), lateral orbital cortex (lORB), retrosplenial cortex (RSC), posterior-anterior part of insular cortex (paIC), anterior cingulate cortex (ACC), and dorsal anterior part of insular cortex (daIC)) and thalamic nuclei (AM, VM, CM, MD, lateral dorsal nucleus (LD), VAL, ventral posterior nucleus (VP), and VPM) (Fig. 4B). Interestingly, higher-dose ketamine did not augment the hypothalamic area except for ARH. These results indicate that the hypothalamic area is the key region affected by lower-dose ketamine, and that the cerebral cortex is a key region affected by higher-dose ketamine.

**Figure 4.**
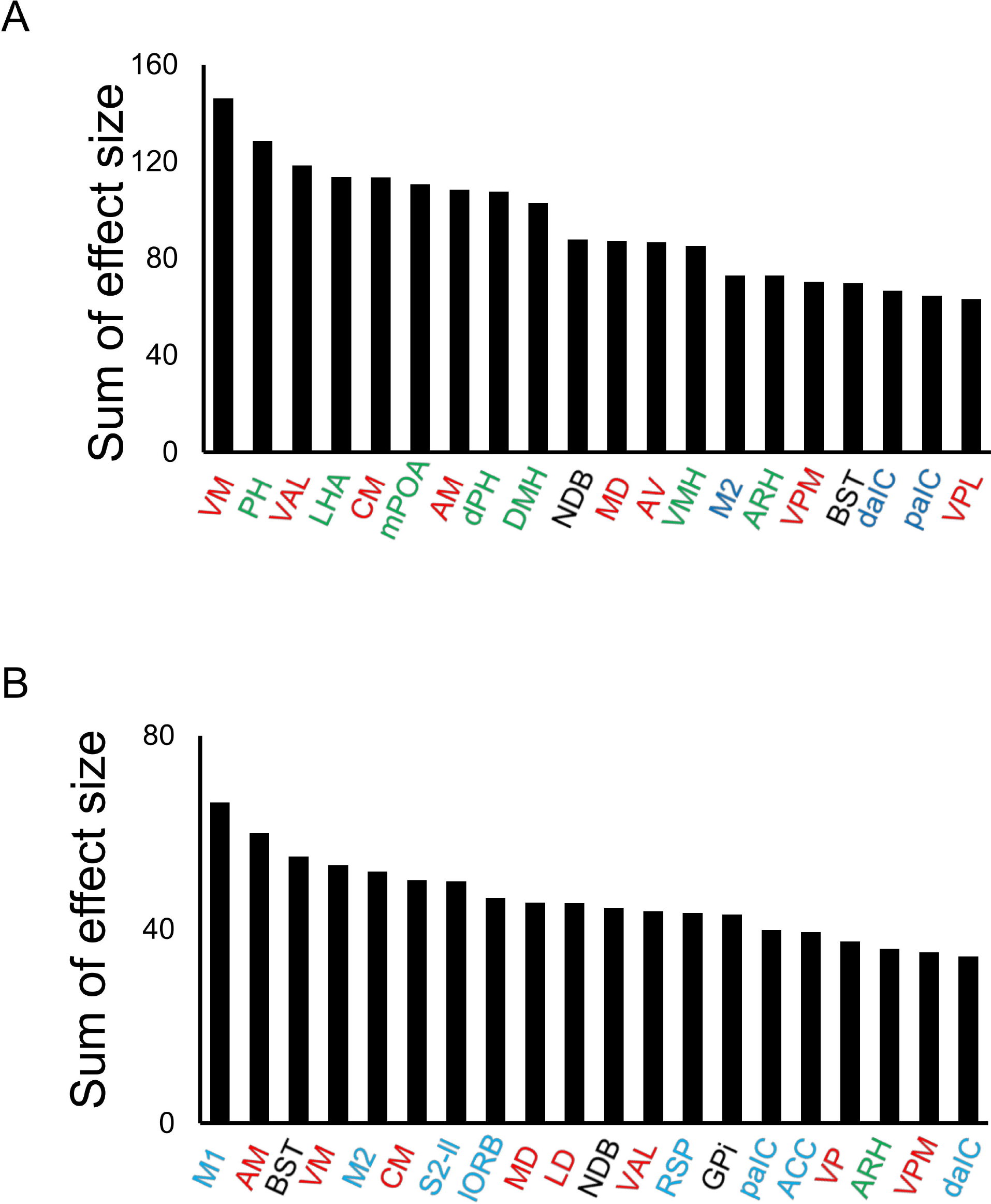
Dose-dependent increase of FC following ketamine injection Extent to which individual regions are affected by the network of strengthened connectivity following (A) lower-dose ketamine and (B) higher-dose ketamine injection. The sum of effect sizes was calculated across all connections associated with a given region. Red, green, and blue regions indicate the thalamic nuclei, hypothalamic nuclei, and the cerebral cortex respectively. AM, anteromedial nucleus; ARH, arcuate nucleus; AV, anteroventral nucleus; BST, bed nuclei of the stria terminalis; CM, centromedial nucleus of the thalamus; DMH, dorsomedial hypothalamus; daIC, dorsal anterior part of insular cortex; paIC, posterior-anterior part of insular cortex; LHA, lateral hypothalamic area; M1, primary motor cortex; M2, secondary motor cortex; MD, mediodorsal nucleus; NDB, nucleus of the diagonal band; (d)PH, (dorsal) posterior hypothalamus; mPOA, medial preoptic area; VAL, ventral anterior-lateral complex; VMH, ventromedial hypothalamic nucleus; VPL, ventral posterolateral nucleus; VPM, ventral posteromedial nucleus.

### 3.3 Graph theory

We used the graph theory to investigate the alterations in FC architecture induced by ketamine injection. The MCC and Eglob significantly increased following lower-dose ketamine injection (p < 0.05) (Figs. 4A and 4B). Lower-dose ketamine also increased Eloc including the thalamus (bilateral VM, bilateral MD, left CM and left VPM) and cerebral cortex (left low-limb of primary somatosensory cortex (SSp-ul), right primary and secondary motor cortex (M1 and M2) and sum of degrees in the right dPH (p < 0.005, uncorrected) (Figs. 4C and 4D). Higher-dose ketamine significantly increased Eglob (p < 0.05), but not MCC (Figs. 4E and 4F). Higher-dose ketamine also increased Eloc in the RSC, sum of degrees in the right RSC and right posteromedial primary visual cortex (V1pm) respectively (p < 0.005, uncorrected) (Figs. 4G and 4H).

## 4. Discussion

In this study, we investigated the dose-dependent alterations in FC following ketamine injection in awake mice. A lower dose of ketamine (5 mg/kg) increased the FC strength in several regions, including the hypothalamic nuclei, thalamic nuclei, and hippocampus. In contrast, a higher dose of ketamine (50 mg/kg) increased the FC strength in several regions, including the thalamus and cerebral cortex. Furthermore, graph analysis showed that the key brain regions with altered FC differed between the lower and higher doses of ketamine.

### 4.1 Whole brain FC in awake mice

Most previous studies counted the c-Fos-positive neurons to investigate altered neuronal activity following ketamine injection. However, this approach should use different animals in the control group, such as the vehicle injection, and ketamine groups. As the resting-state FC architecture has individual variability, a direct comparison of ketamine administration could directly demonstrate the effects of ketamine on FC in the same animal, without considering individual variations confounding the results. Furthermore, an advantage of fMRI is that whole-brain activity can be investigated at several time points, such as before and after ketamine injection. In contrast, c-Fos immunostaining requires an enormous amount of time to investigate the entire brain at several time point. Therefore, based on the hypothesis, most of c-Fos analysis have focused on target brain regions. This could be one reason why most previous studies have investigated the effects of ketamine on the hippocampus, a part of cerebral cortex, and thalamus. Another advantage of fMRI is that FC, which is the dynamic synchronization of the neuronal activity of anatomically different regions, can be investigated. FC includes the temporal dynamics and spatial distribution of functional networks in living animals. Importantly, regional FC changes occur in psychiatric disorders [10,22,23] as well as altered neuronal activity [24].

### 4.2 FC alteration by lower-dose ketamine

Interestingly, the anesthetic dose of ketamine is considered to be more than 100 mg/kg, and it is widely used in combination with xylazine [25], whereas a single injection of ketamine (less than 100 mg/kg) modifies this behavior [26]. Higher-dose ketamine (more than 50 mg/kg) impairs working memory in the Y-maze as well as sensorimotor gating in mice [1]. Furthermore, ambulatory activity is increased by higher-dose ketamine (more than 50 mg/kg). Another study showed that 10 mg/kg ketamine increases hyperlocomotion [27]. Weston et al. summarized the effects of ketamine injection on the behavior of mice [26] and several studies on single-injection ketamine have suggested that a lower dose of ketamine injection, such as 10 mg/kg or less, increases locomotion [28]. Importantly, 10–30 mg/kg ketamine injection is effective in reducing immobility in response to acute stressors, such as the forced swim test [25]. In this study, we found that the head motion increased following lower-dose ketamine injection but the respiration was not changed (Fig. 1C). This can be explained by the increased locomotion. Forced swimming induces c-Fos expression in the ventrolateral preoptic nucleus and DMH in mice [29,30]; however, c-Fos expression following ketamine injection in FST mice has not been investigated. A lower dose of ketamine also produces fast-acting behavioral antidepressant-like effects in mouse models, which depend on the rapid synthesis of brain-derived neurotrophic factors. Chronic mild stress decreases the FC between the hippocampus and thalamus [31]. Previous studies have also revealed that a lower dose of ketamine (30 mg/kg) induces food intake in activity-based anorexia mice [3,32], but there have not been any reports on higher doses. The present study demonstrated that a lower dose of ketamine increased the FC mostly in the hypothalamic nuclei, thalamic nuclei, and hippocampus (Fig. 2). In addition, the sum of the effect sizes in each node showed that the FC with the DMH, medial preoptic area (mPOA), and lateral hypothalamus (LH) increased (Fig. 4A). The graph analysis revealed a significant increase in connections with the dPH and a significant increase in Eloc in several brain regions, including the thalamic and hypothalamic nuclei (Fig. 5A). The significant increase in the MCC indicates a change into a “small-world-network,” meaning dense, tight-knit local neighborhoods [33]. The increased FC among these regions could possibly interact with the decrease in FC owing to acute/chronic stress, resulting in the rescue of abnormal behavior. Altered FC with the hypothalamus is not widely understood because few studies have focused on the relationship between FC and the hypothalamus in stress models and anxiety-based anorexia. Further study of FC in the hypothalamus should be performed in the future.

**Figure 5.**
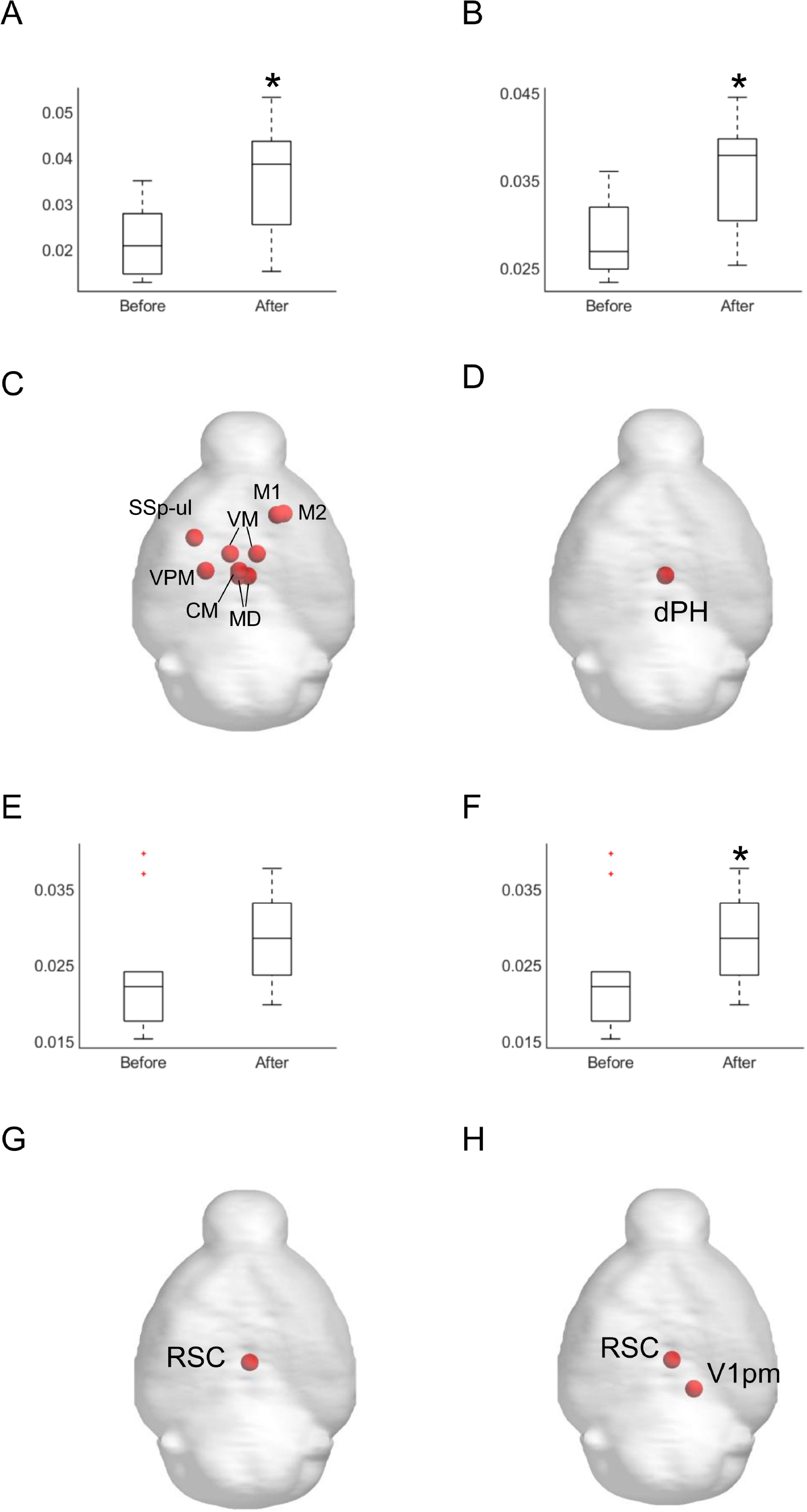
Graph theory (A-D) Significant changes in (A) mean clustering coefficients (p < 0.05), (B) global efficacy (p < 0.05), (C) local efficacy (p < 0.005, uncorrected), and (D) sum of degrees (p < 0.005, uncorrected) following lower-dose ketamine injection. (E-H) Significant changes in (E) mean clustering coefficients (p < 0.05), (F) global efficacy (p < 0.05), (G) local efficacy (p < 0.005, uncorrected), and (H) sum of degrees (p < 0.005, uncorrected) following higher-dose ketamine injection. CM, centromedial nucleus of the thalamus; MOp, primary motor cortex; MOs, secondary motor cortex; SSp-ul, primary somatosensory cortex of the upper limb; VM, ventromedial nucleus of the thalamus; VPM, ventral posteromedial nucleus of the thalamus

### 4.3 FC alteration by higher-dose ketamine

A higher dose of ketamine (50–100 mg/kg), which is still sub-anesthetic, induces ataxia, head weaving, and loss of coordinated movement [2]. Furthermore, more than 50 mg/kg ketamine injection impairs working memory in the Y-maze and sensorimotor gating, as assessed by the inhibition of the acoustic startle response in mice. A 100 mg/kg ketamine injection induces the widespread c-Fos expression throughout the brain in mice [34]. Another study revealed that the injection of 50 mg/kg ketamine increases c-Fos expression in the cingulate and RSC [2]. The present study showed a significant increase in FC across wide regions of the brain. Remarkably, the sums of the effect size showed that the FC in the ACC, RSC, and olfactory bulb was large following the higher ketamine injection, but not following the lower ketamine injection (Fig. 4). In addition, the graph analysis showed a significant increase in the sum of the degrees in the RSC (Fig. 5). The significant Eloc in the RSC indicates that the nodes in a specific, localized neighborhood become more densely interconnected, allowing for faster and more resilient information exchange, routing, and social communication within a specific cluster [35]. Overall, increased FC with the RSC is essential for FC alteration following higher-dose ketamine injection.

In the present study, using awake mouse fMRI, we clearly showed that sub-anesthetic dose of ketamine induced the dose-dependent effect on functional connectivity. The lower-dose (5 mg/kg) ketamine alters the FC mostly among the thalamus, the hypothalamus, and the hippocampus. In contrast, hither-dose (50 mg/kg) ketamine alters the FC in the widespread brain regions including the cerebral cortex and thalamus, and the RSP is one of the key regions for this alteration.

## Acknowledgements

We thank to Dr. Yasuko Sugase for the discussion and advise of experimental paradigm.

## Data availability statement

The data supporting the reported findings are available from the corresponding author upon reasonable request.

## Author Contributions: CRediT

**Tomokazu Tsurugizawa:** conception, design of the work, the acquisition, analysis, interpretation of data and writing the article. **Aya Takemura:** design of the work and writing the article.

## References

1 Qin, Z., Zhang, L., Zasloff, M. A., Stewart, A. F. R. & Chen, H. H. Ketamine’s schizophrenia-like effects are prevented by targeting PTP1B. Neurobiol Dis 155, 105397, doi:10.1016/j.nbd.2021.105397 (2021).

2 Nishizawa, N. et al. The effect of ketamine isomers on both mice behavioral responses and c-Fos expression in the posterior cingulate and retrosplenial cortices. Brain Res 857, 188–192, doi:10.1016/s0006-8993(99)02426-9 (2000).

3 Chen, Y. W., Sherpa, A. D. & Aoki, C. Single injection of ketamine during mid-adolescence promotes long-lasting resilience to activity-based anorexia of female mice by increasing food intake and attenuating hyperactivity as well as anxiety-like behavior. Int J Eat Disord 51, 1020–1025, doi:10.1002/eat.22937 (2018).

4 Wright, P. W. et al. Functional connectivity structure of cortical calcium dynamics in anesthetized and awake mice. PLoS One 12, e0185759, doi:10.1371/journal.pone.0185759 (2017).

5 Palasz, A. et al. Psilocybin and ketamine affect novel neuropeptides gene expression in the rat hypothalamus. J Psychopharmacol 39, 499–508, doi:10.1177/02698811251330783 (2025).

6 Gao, T. H. et al. Chronic lithium exposure attenuates ketamine-induced mania-like behavior and c-Fos expression in the forebrain of mice. Pharmacol Biochem Behav 202, 173108, doi:10.1016/j.pbb.2021.173108 (2021).

7 Duncan, G. E., Moy, S. S., Knapp, D. J., Mueller, R. A. & Breese, G. R. Metabolic mapping of the rat brain after subanesthetic doses of ketamine: potential relevance to schizophrenia. Brain Res 787, 181–190, doi:10.1016/s0006-8993(97)01390-5 (1998).

8 Friston, K. J. Functional and effective connectivity: a review. Brain connectivity 1, 13–36, doi:10.1089/brain.2011.0008 (2011).

9 Kotoula, V., Webster, T., Stone, J. & Mehta, M. A. Resting-state connectivity studies as a marker of the acute and delayed effects of subanaesthetic ketamine administration in healthy and depressed individuals: A systematic review. Brain Neurosci Adv 5, 23982128211055426, doi:10.1177/23982128211055426 (2021).

10 Tsurugizawa, T. et al. Awake functional MRI detects neural circuit dysfunction in a mouse model of autism. Science advances 6, eaav4520, doi:10.1126/sciadv.aav4520 (2020).

11 Mandino, F., Vujic, S., Grandjean, J. & Lake, E. M. R. Where do we stand on fMRI in awake mice? Cereb Cortex 34, doi:10.1093/cercor/bhad478 (2024).

12 Kawazoe, K. et al. Dose-dependent effects of esketamine on brain activity in awake mice: A BOLD phMRI study. Pharmacol Res Perspect 10, e01035, doi:10.1002/prp2.1035 (2022).

13 Tsurugizawa, T. & Yoshimaru, D. Impact of anesthesia on static and dynamic functional connectivity in mice. Neuroimage 241, 118413, doi:10.1016/j.neuroimage.2021.118413 (2021).

14 Bukhari, Q., Schroeter, A., Cole, D. M. & Rudin, M. Resting State fMRI in Mice Reveals Anesthesia Specific Signatures of Brain Functional Networks and Their Interactions. Frontiers in neural circuits 11, 5, doi:10.3389/fncir.2017.00005 (2017).

15 Tsurugizawa, T., Tamada, K., Debacker, C., Zalesky, A. & Takumi, T. Cranioplastic Surgery and Acclimation Training for Awake Mouse fMRI. Bio-protocol 11, e3972, doi:10.21769/BioProtoc.3972 (2021).

16 Power, J. D., Barnes, K. A., Snyder, A. Z., Schlaggar, B. L. & Petersen, S. E. Spurious but systematic correlations in functional connectivity MRI networks arise from subject motion. Neuroimage 59, 2142–2154, doi:10.1016/j.neuroimage.2011.10.018 (2012).

17 Whitfield-Gabrieli, S. & Nieto-Castanon, A. Conn: a functional connectivity toolbox for correlated and anticorrelated brain networks. Brain connectivity 2, 125–141, doi:10.1089/brain.2012.0073 (2012).

18 Rubinov, M. & Sporns, O. Complex network measures of brain connectivity: uses and interpretations. Neuroimage 52, 1059–1069, doi:10.1016/j.neuroimage.2009.10.003 (2010).

19 Onnela, J. P., Saramaki, J., Kertesz, J. & Kaski, K. Intensity and coherence of motifs in weighted complex networks. Physical review. E, Statistical, nonlinear, and soft matter physics 71, 065103, doi:10.1103/PhysRevE.71.065103 (2005).

20 Scharwachter, L., Schmitt, F. J., Pallast, N., Fink, G. R. & Aswendt, M. Network analysis of neuroimaging in mice. Neuroimage 253, 119110, doi:10.1016/j.neuroimage.2022.119110 (2022).

21 Grandjean, J. et al. Common functional networks in the mouse brain revealed by multi-centre resting-state fMRI analysis. Neuroimage 205, 116278, doi:10.1016/j.neuroimage.2019.116278 (2020).

22 Sigurdsson, T., Stark, K. L., Karayiorgou, M., Gogos, J. A. & Gordon, J. A. Impaired hippocampal-prefrontal synchrony in a genetic mouse model of schizophrenia. Nature 464, 763–767, doi:10.1038/nature08855 (2010).

23 Zerbi, V. et al. Brain mapping across 16 autism mouse models reveals a spectrum of functional connectivity subtypes. Molecular psychiatry 26, 7610–7620, doi:10.1038/s41380-021-01245-4 (2021).

24 Oyarzabal, E. A. et al. Chemogenetic stimulation of tonic locus coeruleus activity strengthens the default mode network. Science advances 8, eabm9898, doi:10.1126/sciadv.abm9898 (2022).

25 Zanos, P. et al. NMDAR inhibition-independent antidepressant actions of ketamine metabolites. Nature 533, 481–486, doi:10.1038/nature17998 (2016).

26 Weston, R. G., Fitzgerald, P. J. & Watson, B. O. Repeated Dosing of Ketamine in the Forced Swim Test: Are Multiple Shots Better Than One? Frontiers in psychiatry 12, 659052,

27. doi:10.3389/fpsyt.2021.659052 (2021).

27 Pomrenze, M. B. et al. Ketamine Evokes Acute Behavioral Effects Via mu Opioid Receptor-Expressing Neurons of the Central Amygdala. Biological psychiatry 98, 538–548, doi:10.1016/j.biopsych.2025.04.020 (2025).

28 Polis, A. J., Fitzgerald, P. J., Hale, P. J. & Watson, B. O. Rodent ketamine depression-related research: Finding patterns in a literature of variability. Behavioural brain research 376, 112153, doi:10.1016/j.bbr.2019.112153 (2019).

29 Yanagida, S. et al. Effect of acute imipramine administration on the pattern of forced swim-induced c-Fos expression in the mouse brain. Neurosci Lett 629, 119–124, doi:10.1016/j.neulet.2016.06.059 (2016).

30 Hiraoka, K. et al. Pattern of c-Fos expression induced by tail suspension test in the mouse brain. Heliyon 3, e00316, doi:10.1016/j.heliyon.2017.e00316 (2017).

31 Zeng, T. et al. Nrf2 regulates iron-dependent hippocampal synapses and functional connectivity damage in depression. J Neuroinflammation 20, 212, doi:10.1186/s12974-023-02875-x (2023).

32 Goodwin-Groen, S., Dong, Y. & Aoki, C. Three daily intraperitoneal injections of sub-anesthetic ketamine ameliorate activity-based anorexia vulnerability of adult female mice. Int J Eat Disord 57, 1447–1464, doi:10.1002/eat.24036 (2024).

33 Masuda, N., Sakaki, M., Ezaki, T. & Watanabe, T. Clustering Coefficients for Correlation Networks. Frontiers in neuroinformatics 12, 7, doi:10.3389/fninf.2018.00007 (2018).

34 Hu, Y. et al. Comparative brain-wide mapping of ketamine- and isoflurane-activated nuclei and functional networks in the mouse brain. eLife 12, doi:10.7554/eLife.88420 (2024).

35 Latora, V. & Marchiori, M. Efficient behavior of small-world networks. Phys Rev Lett 87, 198701, doi:10.1103/PhysRevLett.87.198701 (2001).

